# USAG1 protein: An important drug target in teeth regeneration

**DOI:** 10.1101/2022.08.01.502414

**Authors:** R Poorani, E Elakkiya, Krishna Kant Gupta

## Abstract

The BMP antagonist and Wnt signaling modulator, USAG-1 (uterine sensitization associated gene-1) is mostly expressed in the kidneys and suppresses BMP7 bioactivity, which is crucial for tooth development. The homodimeric 35-kDa protein known as bone morphogenetic protein-7 is essential for the specification and patterning of the early embryo and controls apoptosis in a variety of developmental events. A single-gene knockout (KO) mouse model for Usag-1, also known as Sclerostin domain containing 1 (SOSTDC1), ectodin, Wnt modulator in surface ectoderm (WISE), CCAAT/enhancer-binding protein beta (CEBPB), Sprouty homolog 2 (SPRY2), Sprouty homolog 3 (SPRY3), or Epiprofin (EPFN), has demonstrated that Usag1 inhibition is important for teeth regeneration in mice. Anti Usag1 method was done by other scientists outside India. Tideglusib when placed at damaged teeth, it increases WNT signaling by inhibiting GSK-3.It results into teeth repair .Suginami et al proposed USAG-1 as a promising drug target for teeth regeneration .They proposed anti-USAG-1 approach to induce teeth regeneration but it has been associated with many sideaffects. This study aims to provide a natural lead molecule as a therapeutic drug for teeth regeneration.

## 1. INTRODUCTION

One of the vertebrate organs whose molecular development is being investigated is the tooth. The mediating role of secretory signaling molecules in facilitating sequential and reciprocal inductive connections between the oral epithelium and mesenchyme has already been established. Finally, a signaling or organising centre that expresses the same signals as other organising centres in the embryo and appears to constrain tooth structure was discovered in the dental enamel knot. The molecules from the bone morphogenetic protein (BMP) and Wnt that control the morphogenesis of a single tooth are examples of molecular signals that regulate organogenesis. BMP signaling is necessary for the morphogenesis of extra teeth [4], whereas Wnt signaling is crucial for the development of extra teeth [5]. It is unclear, though, whether BMP or Wnt signaling is necessary to count the teeth. A person’s total number of teeth varies from person to person. Between 0.1 and 3.6 percent of people can have congenital increases or decreases [6]. Supernumerary teeth, often known as teeth agenesis, refer to any orthodontic structure that is either more or less numerous than the average number of teeth in a person.

**Fig 1.1:**
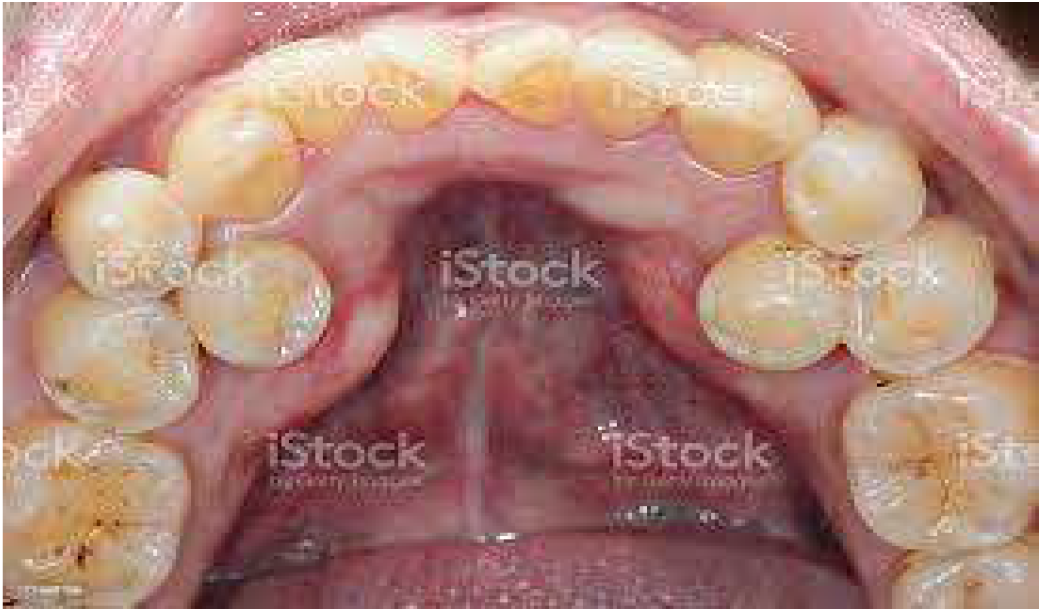
Supernumerary teeth condition in human

**Fig 1.2:**
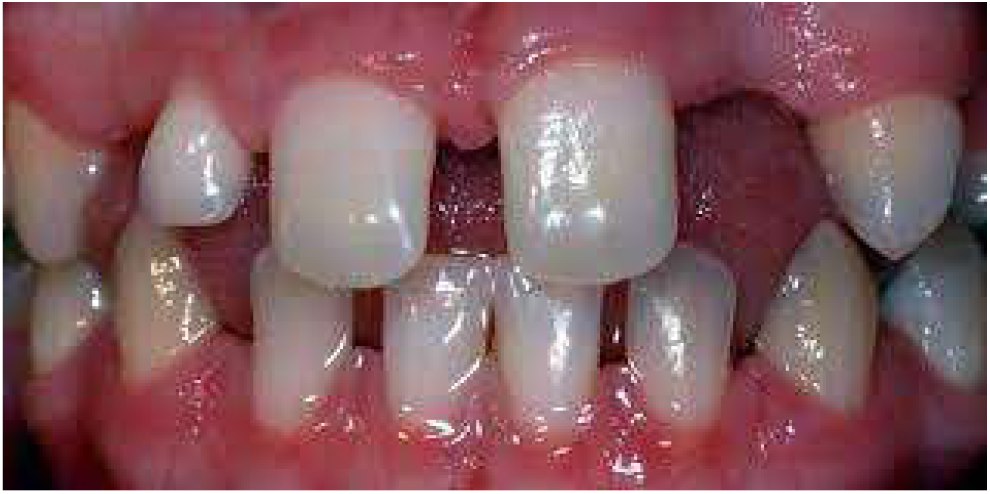
Teeth agenesis condition in human

A BMP antagonist and Wnt signaling modulator, USAG-1 (uterine sensitization associated gene-1) is mostly expressed in the kidneys and suppresses BMP7 bioactivity, which is crucial for tooth development [1]. The homodimeric 35-kDa protein known as bone morphogenetic protein-7 is essential for the specification and patterning of the early embryo and controls apoptosis in a variety of developmental events [2,3]. A single-gene knockout (KO) mouse model for Usag-1, also known as Sclerostin domain containing 1 (SOSTDC1), ectodin, Wnt modulator in surface ectoderm (WISE), CCAAT/enhancer-binding protein beta (CEBPB), Sprouty homolog 2 (SPRY2), Sprouty homolog 3 (SPRY3), or Epiprofin (EPFN), has demonstrated that Usag1 Usag1 inhibition is important for teeth regeneration in mice[7].

**Fig 1.3:**
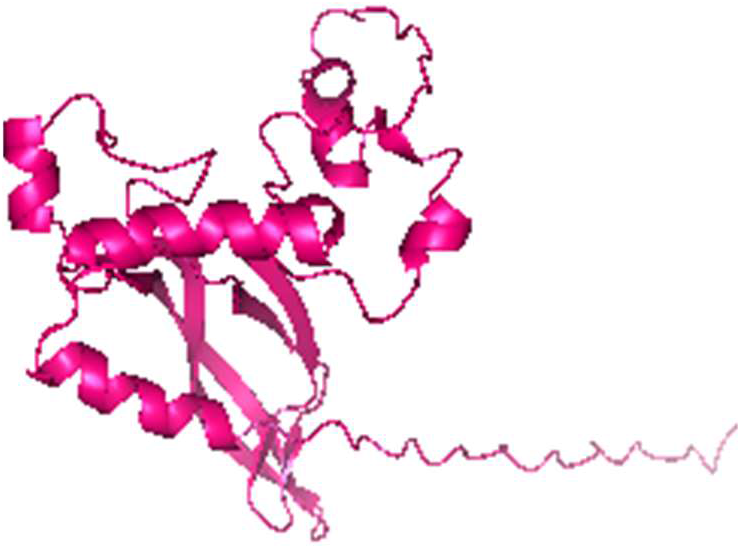
Modeled USAG1 Protein

Supernumerary teeth have been linked to USAG1 deficiency in mice in earlier studies [7]. The progressive development of the rudimentary upper incisor appears to be the cause of the supernumerary maxillary incisor’s formation. BMPs are expressed in the mesenchyme and epithelium of the rudimentary incisor in E14 and E15, according to earlier studies. In the E15 incisor, supernumerary teeth do not develop when BMP signaling is inhibited. Many genes, like Msx1, Runx2, Ectodysplasin A (EDA), or Pax9, have been identified as the cause of oral agenesis using KO mouse models. According to studies, tooth development that had been halted in Runx2/mice, a mouse model for congenital tooth agenesis, might be restored in Runx2/USAG-1/ animals, a supernumerary mouse model. Additional teeth can be treated with targeted molecular therapy for tooth regeneration.

Anti Usag1 method was done by other scientists outside India. Tideglusib when placed at damaged teeth, it increases WNT signaling by inhibiting GSK-3.It results into teeth repair .Suginami et al proposed USAG-1 as a promising drug target for teeth regeneration .They proposed anti-USAG-1 approach to induce teeth regeneration but it has been associated with many side-affects.

**Fig 1.4:**
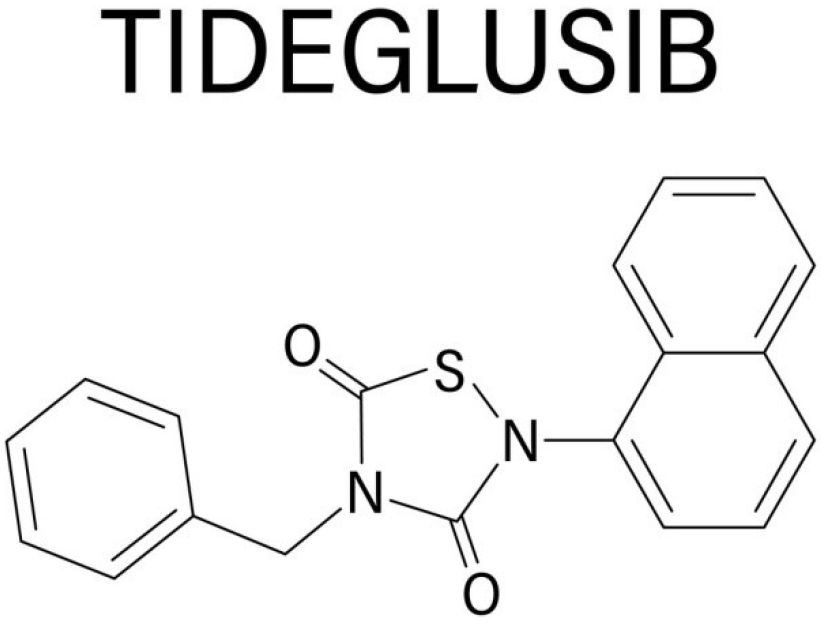
Tideglusib compound

With the help of molecular dynamic simulation, free binding energy calculations, and structurally based virtual screening, this study aims to create a natural anti-USAG1 antibody molecule for local tooth development arrest and recovery.

Previous international research was conducted in Japan based on

1. Anti-USAG-1 therapy for increased BMP signaling in tooth regeneration.
2. Using targeted molecular therapy to regulate tooth loss, the development of tooth regenerative medicine techniques.
3. Due to USAG-1 abrogation, rudimentary incisors persist and erupt as extra teeth.

This study aims to provide a natural lead molecule as a therapeutic drug for teeth regeneration.

## 2. METHODOLOGY

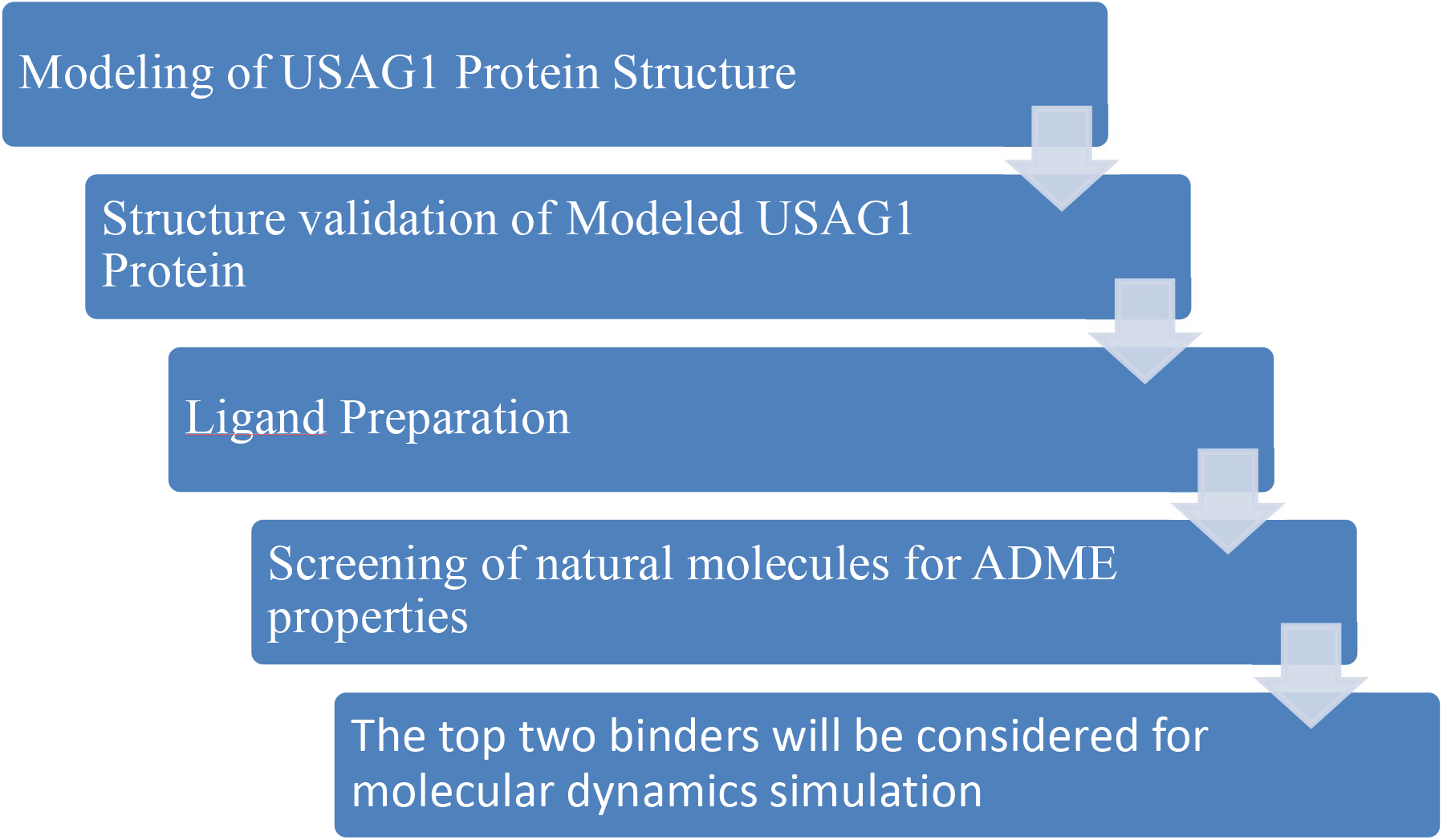

### 2.1 Model building and its validation

The protein data library did not contain the USAG1 protein’s structural structure. The USAG1 protein’s raw sequences were discovered in Uniprot (Q6X4U4_1). So, the usag1 protein’s FASTA sequences were obtained. It was blasted with protein blast and a comparable structure to the usag1 protein was discovered. The templates for the protein (2k8p A,6L6R C,2KD3) and comparable structures of the usag1protein were found. The highest similarity score was used to select the templates, and 2k8p A obtained the highest score. Therefore, the usag1 protein structure was modelled using this as a template, and the missing residues were filled using modeller software and by three additional servers: phyre, I-tasser, and Robetta. A technology called MODELLER uses three-dimensional protein models to create. After receiving an alignment of a sequence to be modelled with recognised complexes, MODELLER automatically creates a model incorporating all non-hydrogen atoms[9]. A set of online resources called Phyre2 is available to forecast and examine changes in protein structure, function, and mutations[10]. I-TASSER was initially developed for simulations of iterative threading and assembly used in modelling protein structure. By comparing structure predictions with recognised functional templates, it was recently expanded for structure-based function annotation[11]. Robetta is an online programme that provides automated structure prediction and analysis tools for extrapolating protein structural information from genetic data. [12]. This application makes use of ROSETTA, which integrates the Monte Carlo method while taking into account a space of conformations, to find a minimal energy conformation. By changing the torsion angles between a fragment in an acquired model and known structure fragments, Rosetta can study structure. The process produced five models for each protein. [15].

### 2.2 STRUCTURE VALIDATION OF THE MODELLED STRUCTURES

The modeled structures which were retrieved from protein modeling servers(modeler, phyre, I-tasser, Rodbetta) were validated by using the Savesv6.0 server. The SAVES v6. 0 toolkit’s five tools—ERRAT, VERIFY3D, PROVE, PROCHECK, and WHAT CHECK—predict various types of stereochemical features of the protein structure. The structural stereochemical property is predicted by the Ramachandran plot. [13]. The quality score was not good in any of the models. To refine the low quality models,3D refiner and Galaxy web was used. The refined structures were again validated by using Savesv6.0.Among all the refined models, the best model which had a better quality score was chosen and it was proceeded to docking.

### 2.3 DATABASE SEECTION AND LIGAND PREPARATION

We have utilized four different databases to find out natural molecules a)Antioxidant compounds from Medchem b) Anti-inflammatory compounds from lifechemicals c) coconut database and d)skellchem database. The whole compounds sdf file was retrieved from the databases and it was separated into each sdf files using pymol.

### 2.4 STRUCTURE BASED VIRTUAL SCREENING

Docking was done on quality checked usag1 protein with the Anti-inflammatory compounds(2000 compounds) found in lifechemicals database using pyrx software. PyRx is a programme for virtual screening that can be used in computational drug discovery to test libraries of compounds against putative therapeutic targets[14]. A CTCK domain can be found in USAG1 protein (C-terminal cystine knot domain). The Cys-X residue in the knot’s C-terminal cystine knot domain, which are crucial for the CTCK domain to become dimerize, stabilises the protein’s structural integrity. The residues from 75-170 in usag1 protein covers the CTCK domain. Residue number 104 cystine is very important for dimerization and stability of protein so that was considered as active site of binding. USAG1 protein grid box was generated including the residue number 104. After screening, the compounds which had good binding affinity were selected and they were checked for ADME prediction.

### 2.5 ADMET prediction

Absorption, Distribution, Metabolism, Excretion, and Toxicity are collectively referred to as ADMET. Because these characteristics are to blame for over 60% of all medicine failures in clinical trials, predicting ADMET attributes is crucial for the drug design process. Since compounds with poor ADME properties are now removed from the drug development pipeline from the beginning of the process rather than at the end as was the case in the past, significant research and development cost reductions are achieved.

ADME was checked for all the good binding affinity compounds with the help of servers like osiri and SwissADME. Using this the compounds which had good ADME without toxicity was filtered.

The good compounds which had good ADME With good binding affinity was considered for molecular dynamic simulation.

### 2.6 MOLECULAR DYNAMIC SIMULATION

To better understand the molecular stability of simulated three-dimensional complex structures, molecular dynamics simulations were performed using GROMACS 5.0.2 and the GROMOS96 43a1 all-atom force field. The topology of the ligand and protein was designed. The ligand topology has been produced by the PRODRG server. In a cubic box of solution that was 1.50 nm deep, the protein ligands were submerged. All directions made advantage of the periodic boundary system. The solvated systems were neutralised by adding sodium ions in place of the water molecules. The GROMACS software program’s correct features were used to save and analyse the structural coordinates for every 2 ps. Using RMSD, RMSF, H-bond, and Radius of gyration, the least energy complicated structures were compared. It was possible to recover six frames of the complex from the MD trajectory at 0 ns, 10 ns, 20 ns, 30 ns, 40 ns, and 50 ns as PDB files. These frames were then examined to determine which H-bonds stayed constant during the simulation.

## 3. RESULTS AND DISCUSSION

### 3.1 Protein preparation

The usag1 protein structure did not have any structure in protein data bank. So the protein was modeled using software like modeler, rodbetta, I-tasser and phyre. For that the fasta sequences were retrieved from the uniport (Q6X4U4_1).

>sp|Q6X4U4|SOSD1_HUMAN Sclerostin domain-containing protein 1 OS=Homo sapiens OX=9606 GN=SOSTDC1 PE=1 SV=2

MLPPAIHFYLLPLACILMKSCLAFKNDATEILYSHVVKPVPAHPSSNSTLNQARNGGRHF

SNTGLDRNTRVQVGCRELRSTKYISDGQCTSISPLKELVCAGECLPLPVLPNWIGGGYGT

KYWSRRSSQEWRCVNDKTRTQRIQLQCQDGSTRTYKITVVTACKCKRYTRQHNESSHNFE

SMSPAKPVQHHRERKRASKSSKHSMS

After retrieval of the sequences it was blasted using blast-p and got the template id as 2K8P,6L6R, 2KD3 with maximum of 42.37 % as percentage identity.

Using this as template usag1 was modeled using modeler, i-tasser and phyre and the structure was not good and the quality score was less while evaluating the model using saves server.

**Fig 1.1:**
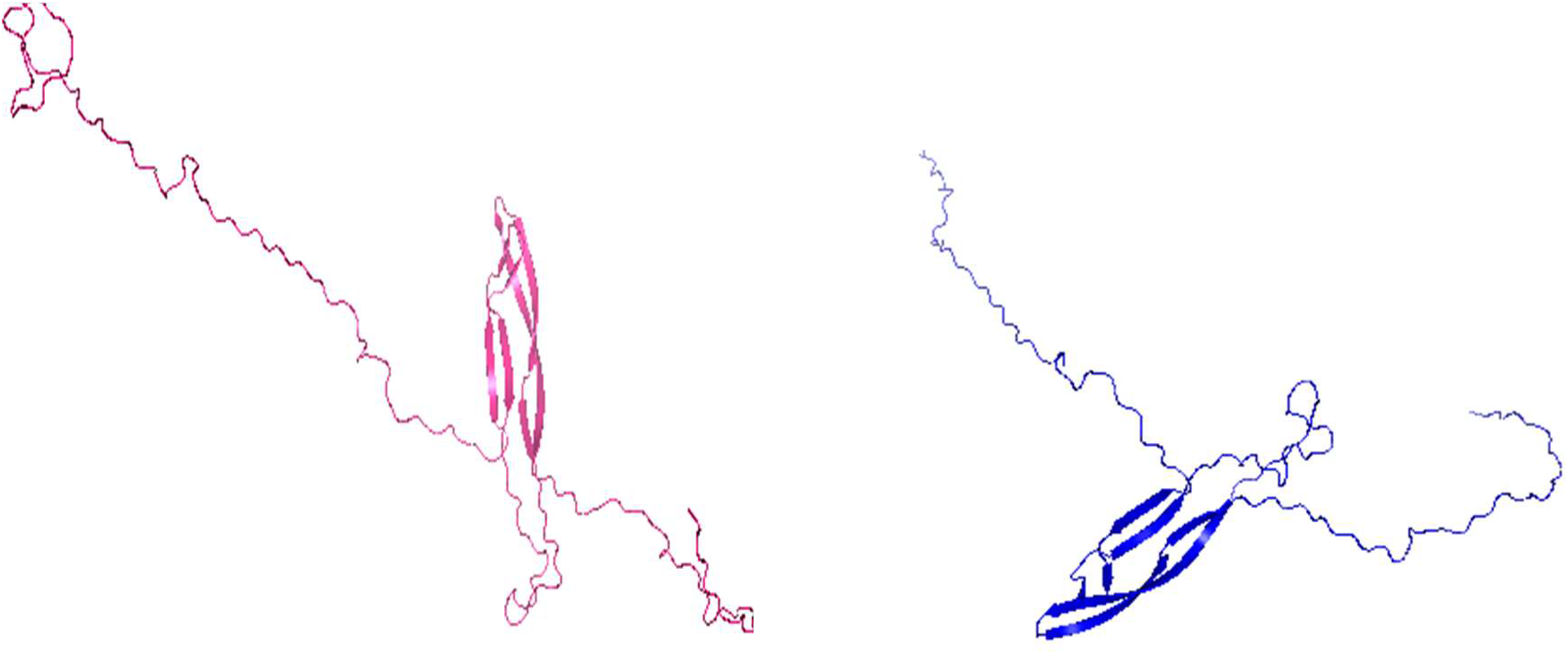
Modeled usag1 protein using modeler and phyre

**Fig 1.2:**
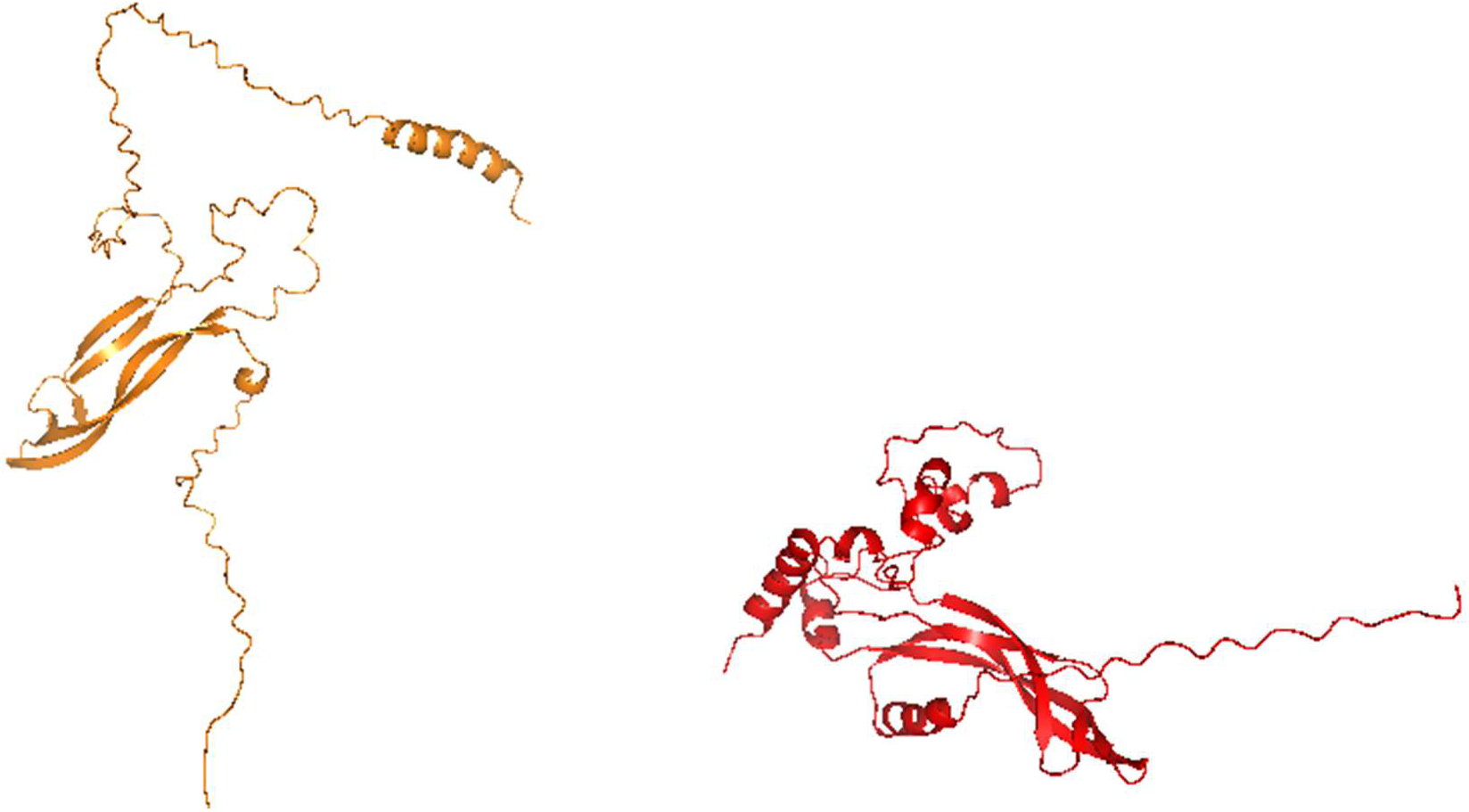
Modeled usag1 protein using google deepmind and rodbetta

Because of the low quality of the modeled structures it was refined using refinement servers like 3D Refiner and Galaxyweb .The best one scored was taken and used that structure for refinement.

**Fig 1.3:**
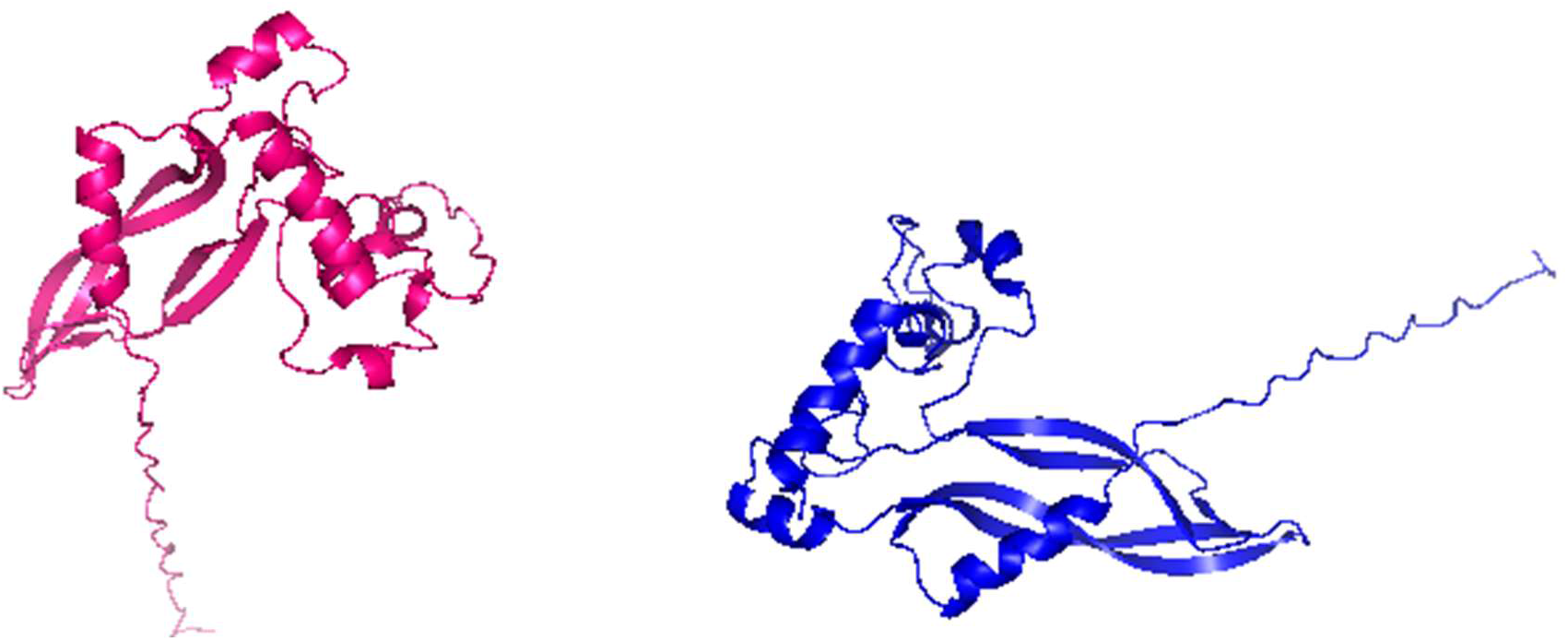
Refined models retrieved from servers

### 3.2 STRUCTURE VALIDATION OF THE REFINED STRUCTURES

Structure validation of refined structures was done using saves errat server.

The quality was good and the best one model among others is the galaxy web server model which had a quality score of 97 which was a good score.

### 3.3 PREPARATION OF LIGAND LIBRARY

Four different databases was used to find out natural molecules a)Antioxidant compounds from Medchem b) Anti-inflammatory compounds from lifechemicals c) coconut database and d)skellchem database.

In this Anti-inflammatory molecules was used because to induce teeth regeneration and to reduce inflammatory in teeth.

### 3.4 STRUCTURE BASED VIRTUAL SCREENING USING PYRX

Virtual screening of the produced ligands was done at the lipase active site. Eight ligands were filtered out of the active site’s 2000 ligands. The cystine at residue number 104 was determined to be the binding site. Table 1.3 displays the top ten ligands according to the binding affinity score in Discovery Studio. Based on the binding affinities in this table, two compounds, 3411 and 0266, have high affinities.

**Table 1.1:** Quality score of the modeled protein from errat server before refinement.

| SERVERS | QUALITY FACTOR |
| --- | --- |
| Modeler | 28 |
| Phyre | 33 |
| I-tasser | 40 |
| Rodbetta | 12 |
| Google Deepmind | 60 |

**Table 1.2:** Quality score of the modeled protein from errat server after refinement.

| SERVERS | QUALITY FACTOR |
| --- | --- |
| GALAXY WEB | 97 |
| 3D refinement | 90 |
| Google deepmind | 76.16 |

**Table 1.3:** screening of natural molecules using pyrx

| Ligand | Binding Affinity | RMSD/ub | RMSD/lb |
| --- | --- | --- | --- |
| model_5_F0266-0248_uff_E=684.47 | -8.3 | 0 | 0 |
| model_5_F3411-4060_uff_E=438.72 | -8.2 | 0 | 0 |
| model_5_F5023-0088_uff_E=682.16 | -8.2 | 0 | 0 |
| model_5_F6473-0874_uff_E=220.55 | -8.1 | 0 | 0 |
| model_5_F0466-0047_uff_E=513.26 | -8.1 | 0 | 0 |
| model_5_F0300-4060_uff_E=438.72 | -8.1 | 8.6 | 5.9 |
| model_5_F0359-0188_uff_E=684.47 | -8.1 | 7.4 | 4.5 |
| model_5_F0758-4060_uff_E=438.72 | -8 | 4.7 | 3.7 |

Screening of reference molecules like tideglusib, 6 Bromoindirubin 3’ oxime, CHIR99021 was done using pyrx. The binding affinity was greater than the previous screened natural compounds which is shown in the table 1.4.

**Table 1.4:** screening of reference molecules using pyrx This proves the natural compounds are better than the already clinically used compounds.

| Ligand | Binding Affinity | RMSD/ub | RMSD/lb |
| --- | --- | --- | --- |
| Tideglusib | -7.9 | 0 | 0 |
| CHIR99021 | -8.2 | 0 | 0 |
| 6 Bromoindirubin 3' oxime | -8.1 | 0 | 0 |

### 3.5 ADMET Prediction of the screened natural compounds

Absorption, Distribution, Metabolism, Excretion, and Toxicity are all acronyms for ADMET.To check whether the drug compounds are eligible for clinical trial and for clinical use without any toxicity, ADME prediction is done.

ADMET prediction is done for all the screened compounds is shown in the table 1.5 and ADME prediction is done for reference compounds is shown in the table 1.6.

**Table 1.5:**
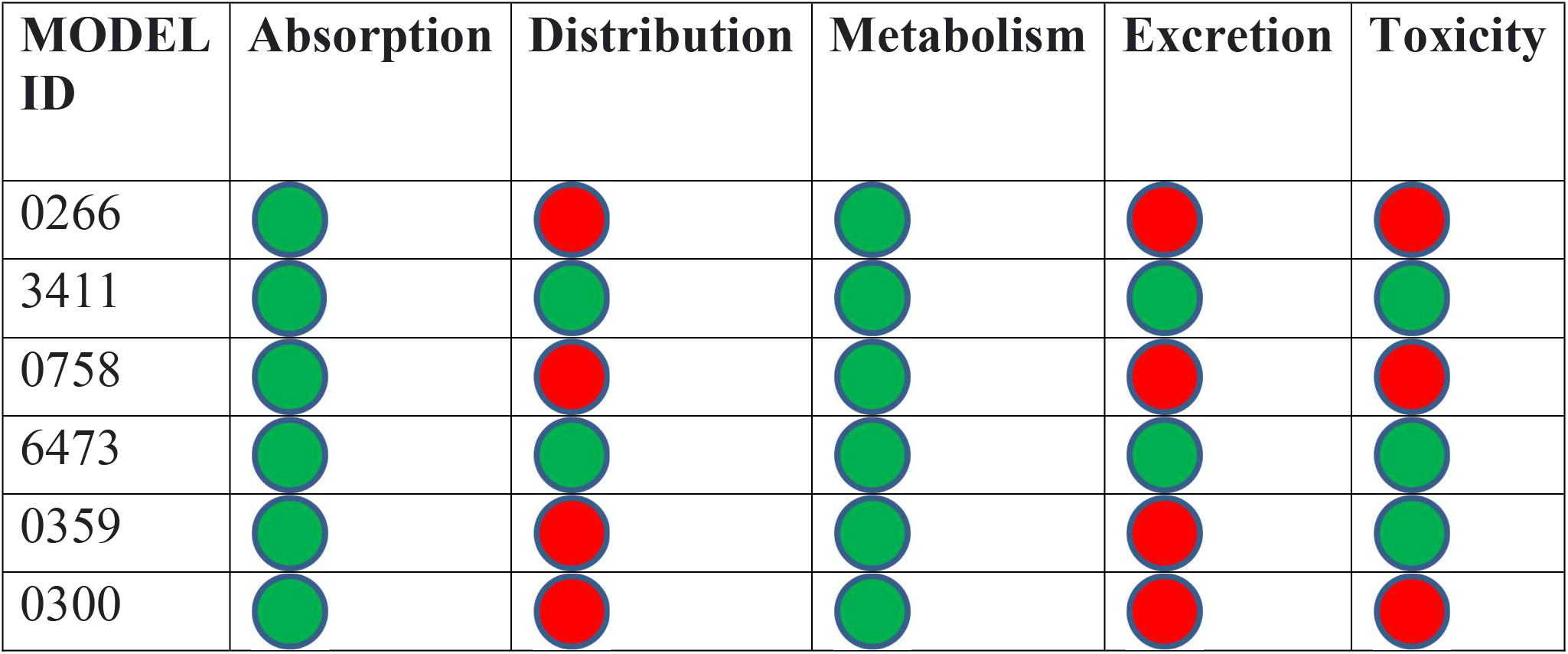
ADMET prediction done for all the screened compounds

**Table 1.5:**
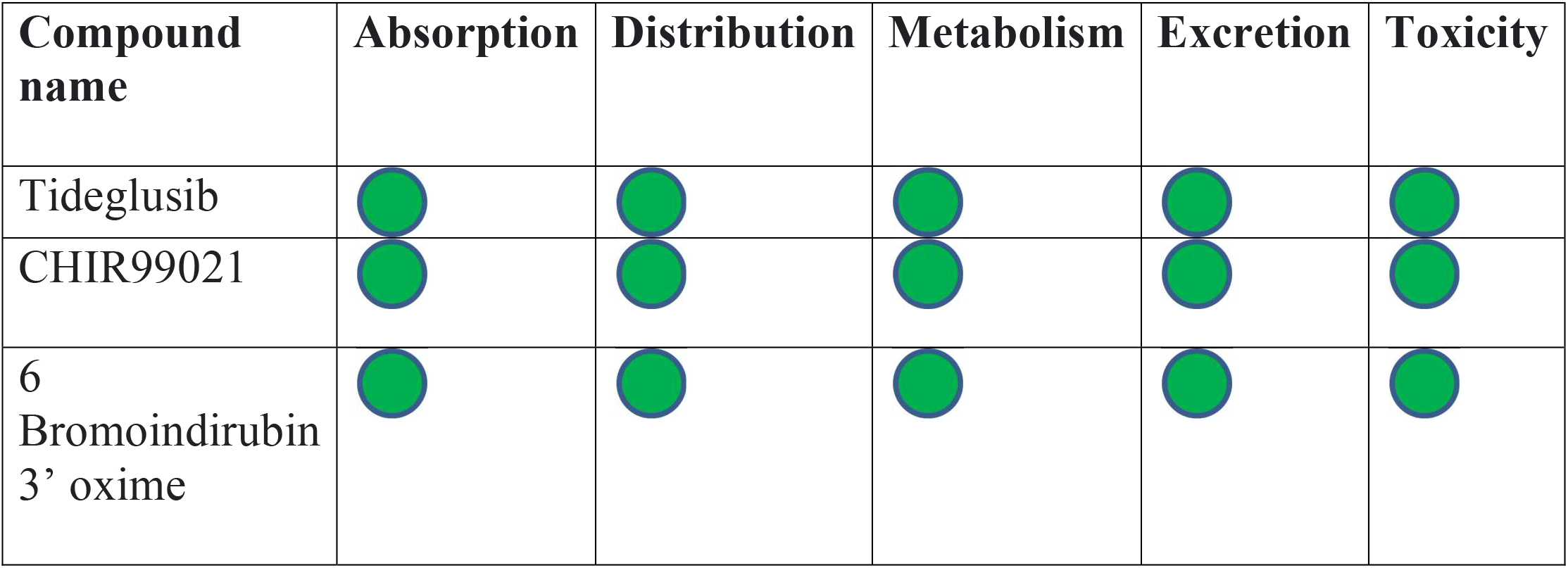
ADMET prediction done for the reference compounds

**Table 1.6:** Mmpbsa analysis for Complex 1,complex 2 and reference compound.

| Ligand name/number | $\Delta E_{vdw}$ (kcal/mol) | $\Delta E_{elec}$(kcal/mol) | $\Delta G_{GB}$(kcal/mol) | $\Delta G_{mmpbsa}$ (kcal/mol) |
| --- | --- | --- | --- | --- |
| 3411 | -29.954 | -5.890 | -21.72 | -10.785 |
| 6473 | -15.869 | -5.869 | -12.52 | -12.52 |
| Tideglusib | -38.37 | -19.146 | -8.72 | -25.69 |

Based on the results it is shown that the compound 3411 and 6473 is good without any toxicity. Though the compound 6473 binding affinity is not good compared to 0266 it is non toxic and has a better moderate binding affinity compared to others. So these compounds 6473 and 3411 can be considered for molecular dynamic simulation.

### 3.6 Ligand interaction of the selected compounds with Usag1 protein

**Fig 1.4.**
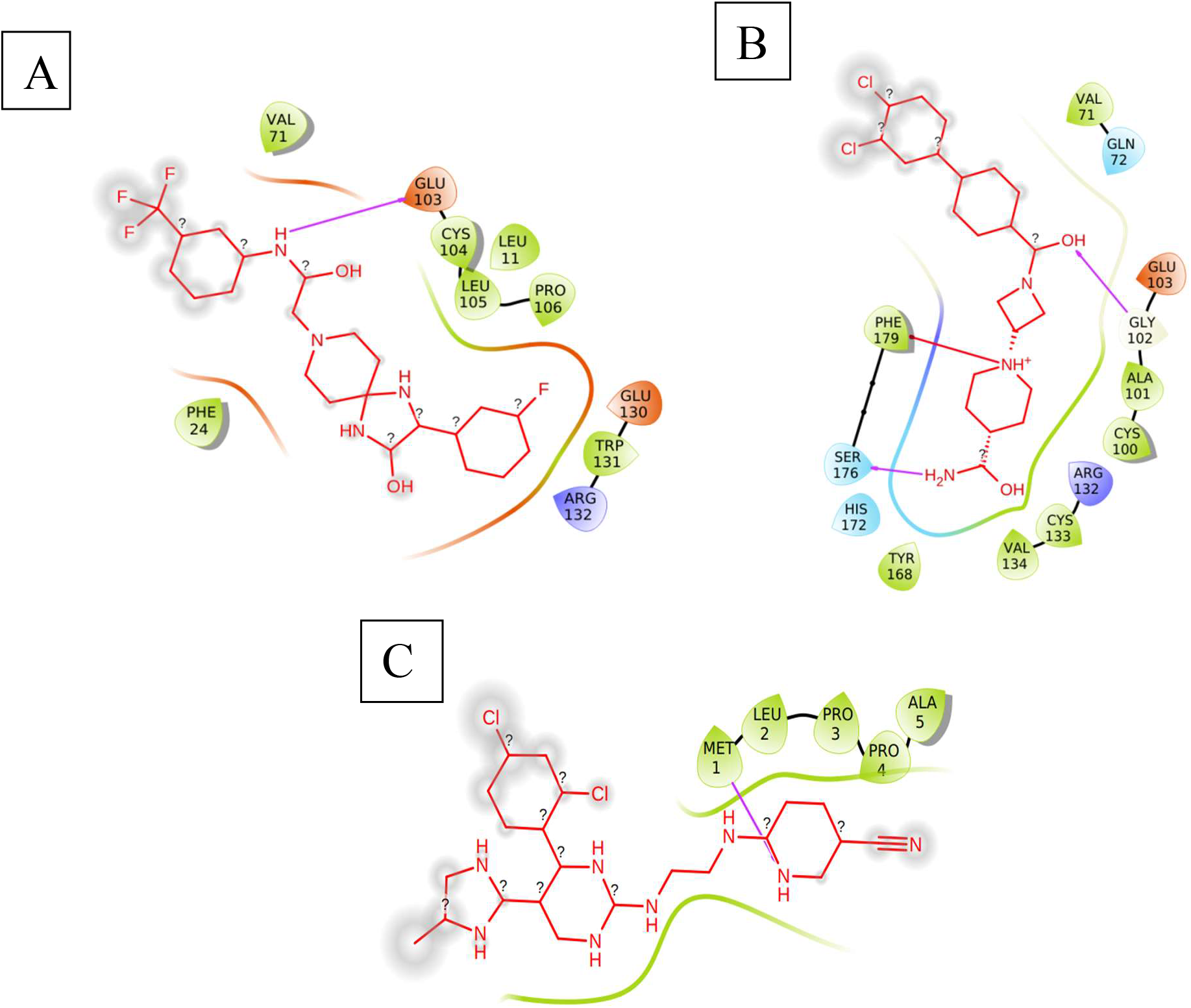
(A): Residues involved in 3411 compound (B): Residues involved in 6473 compound (C): Residues involved in reference compound

#### Hydrogen bond interaction of the selected compounds with Usag1 protein

The lines in the figure indicate the hydrogen bond interaction between the protein and ligand.

**Fig 1.5.**
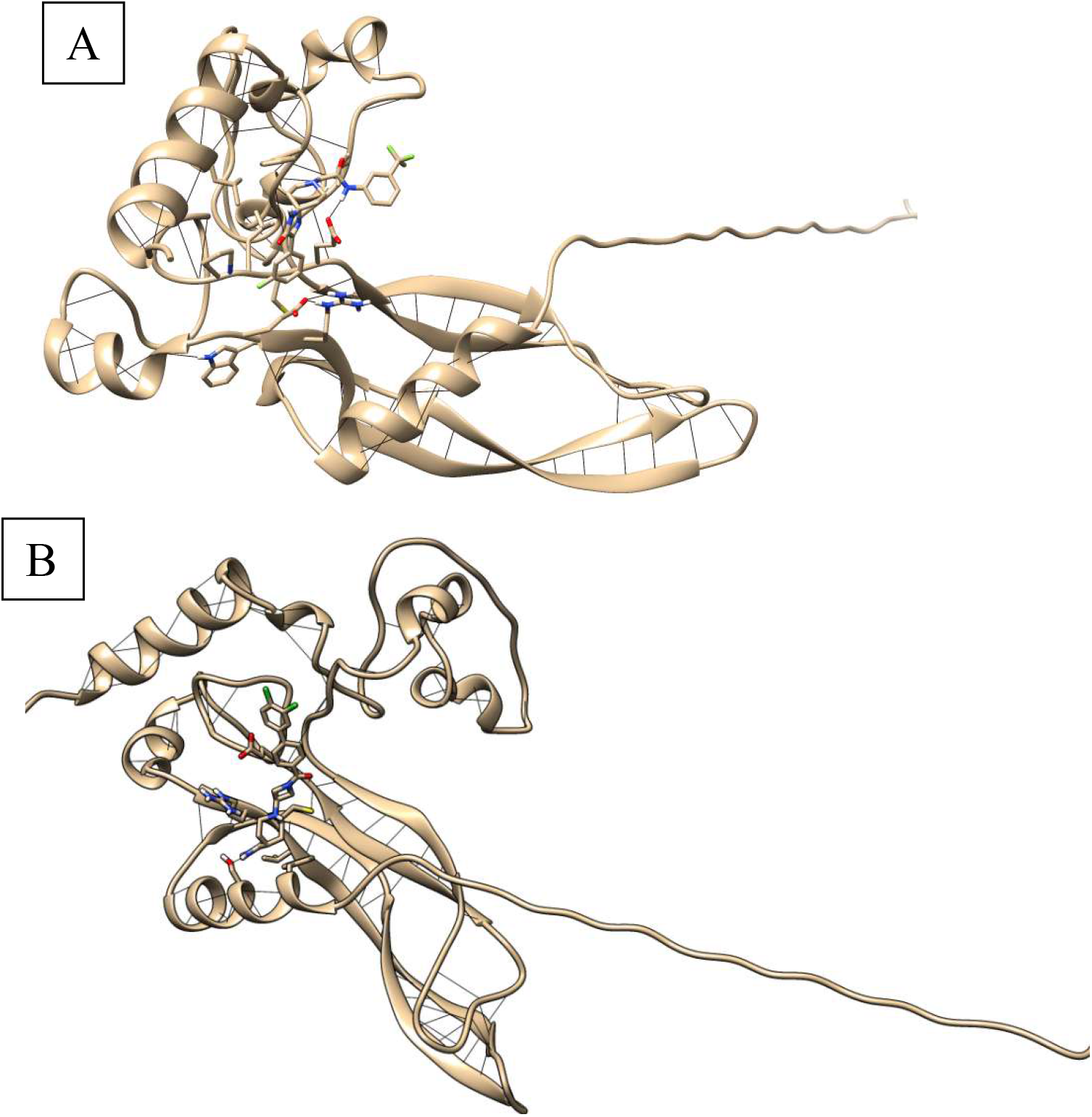
(A): H-bond interaction between the protein and 3411 compound. (B): H-bond interaction between the protein and 6473 compound.

### 3.7 MOLECULAR DYNAMIC SIMULATION

To comprehend and validate the stability of the modeled USAG1 protein structure, the drug compounds which passed the ADME (3411-Complex 1,6473-Complex 2) and the reference compound tideglusib, a 100 ns MD simulation was run. The backbone RMSD profile among all the compounds reveals that the protein may maintain equilibrium until the final production MD. The Complex 2-6473 had a maximum RMS deviation of 1.4 nm, after around 20 ns. Compared to complex 1,complex 2 was stable without much deviations which is shown in the figure 1.4. Additionally, the protein, Complex 1,Complex 2 and reference compound RMSF profile was examined to take into account the structure’s flexible areas . Compared to Complex 1, Complex 2 had maximum fluctuation at 1 ns which falls under (40-50) residue which is shown in the figure 1.5.

**Fig 1.4:**
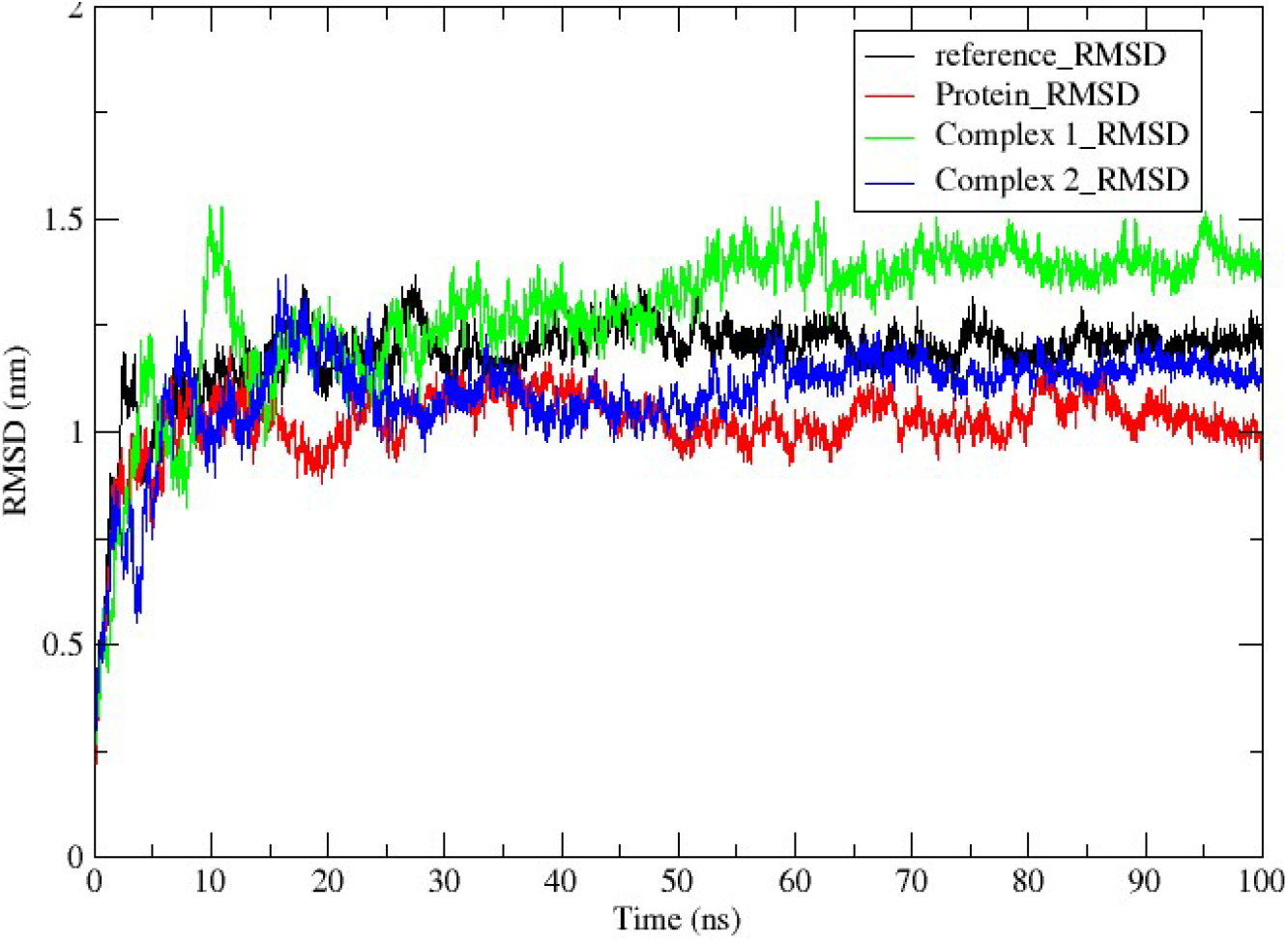
backbone RMSD profile among all the compounds

**Fig 1.5:**
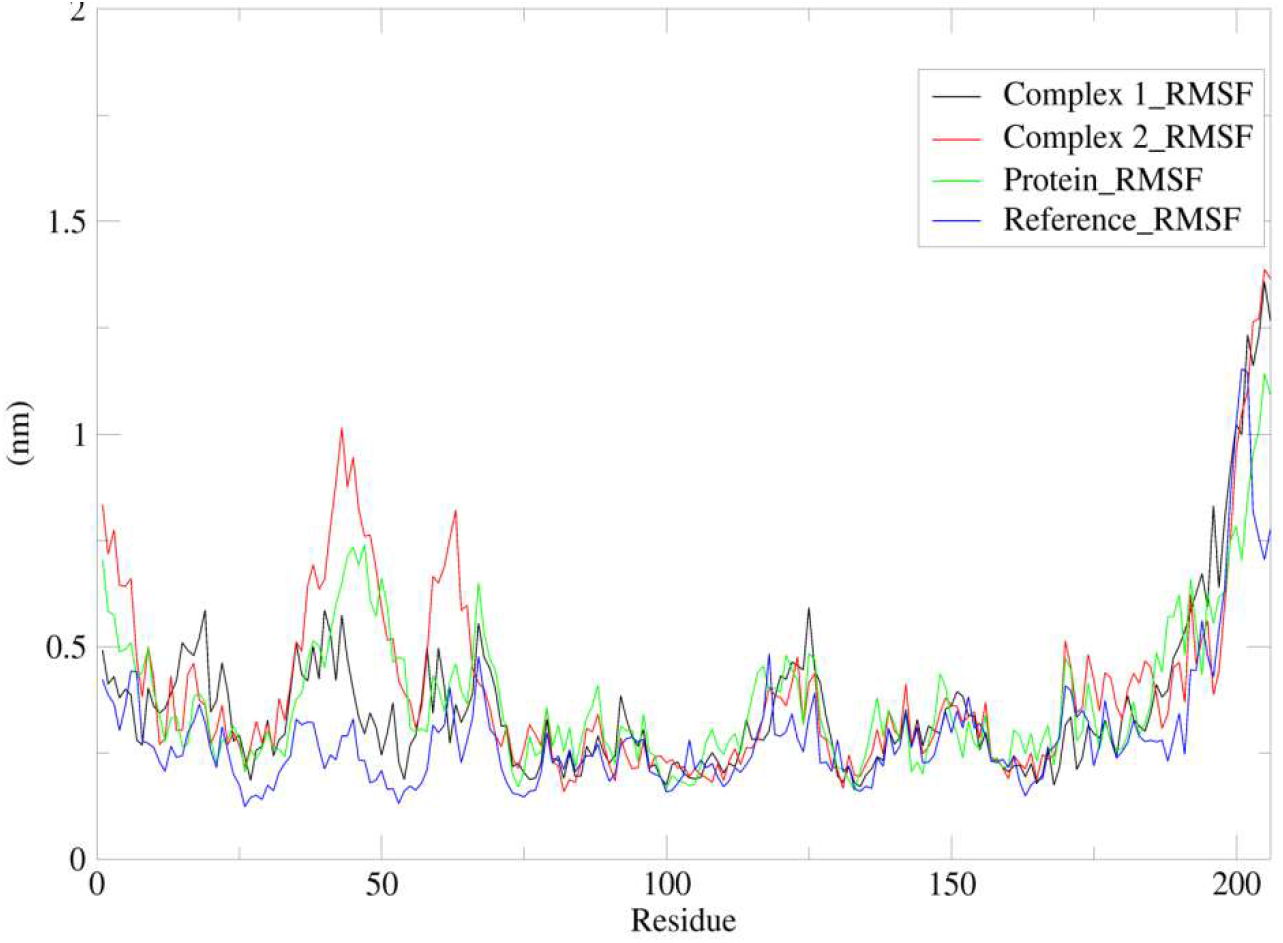
The protein, Complex 1,Complex 2 and reference compound RMSF profile

The radius of gyration was used to determine the protein structure’s compactness; a greater value indicated a less compact structure, while a lower value indicated a more compact one. Complex 2 had lesser value when compared to complex 1 which is shown in the figure 1.6.

**Fig 1.6:**
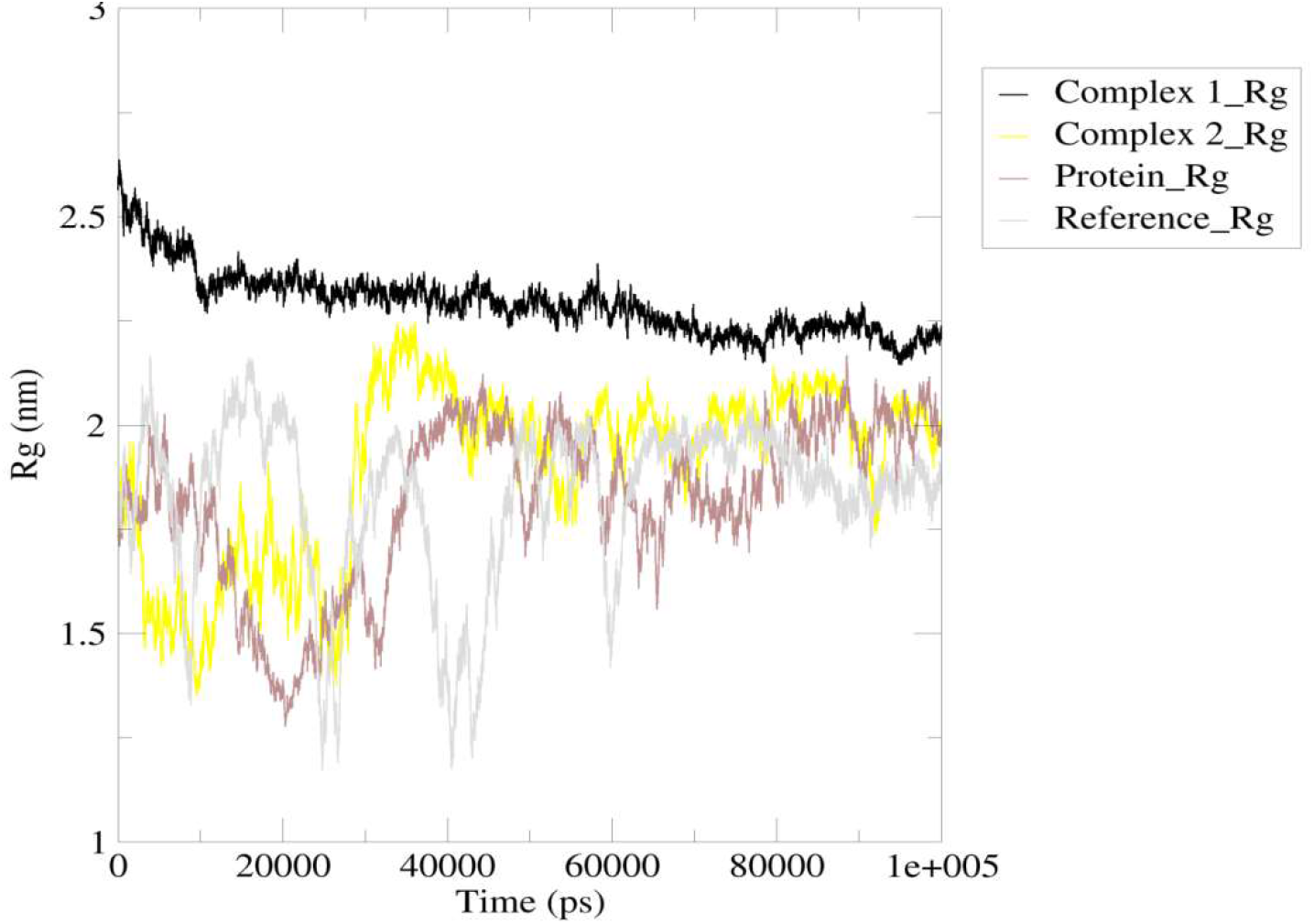
The protein, Complex 1,Complex 2 and reference compound Radius of gyration profile.

It was determined how much of a protein’s amino acid residue is exposed to water, or the solvent accessible surface area (ASA). Given that the active regions of proteins are frequently found on their surfaces, this is a crucial structural characteristic. Comparing Complex 2 to Complex 1, as seen in figure 1.7, Complex 2 has a better exposure.

**Fig 1.7:**
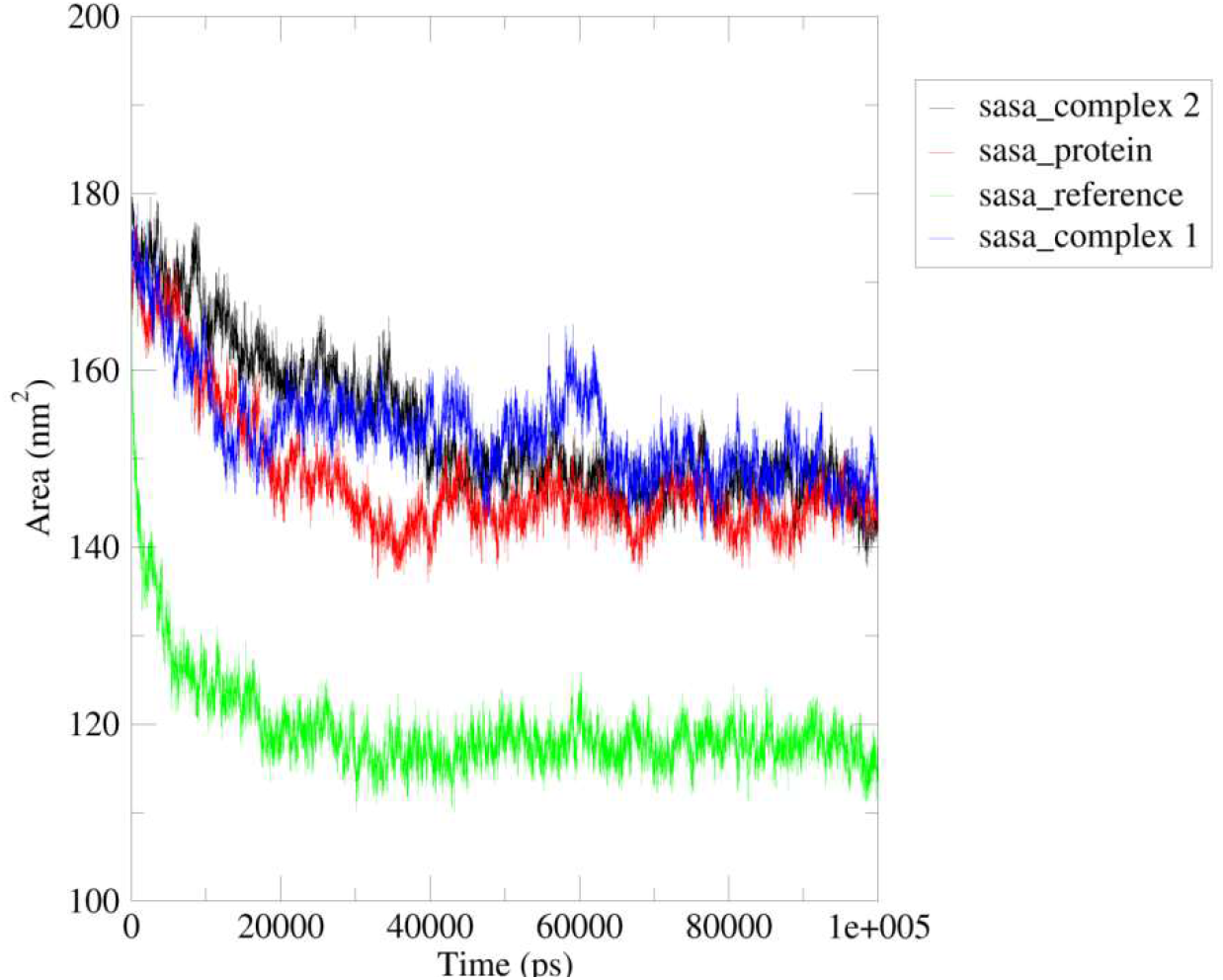
SASA analysis for protein, Complex 1,Complex 2 and reference compound

### 3.8 HYDROGEN BOND

H-Bond was calculated to know the hydrogen interaction between the protein and the ligand which determines the stability.

Complex 2 has a better hydrogen bond interaction with protein and the ligand to the maximum number 6 compared to complex1 which is shown in the figure 1.8.

**Fig 1.8:**
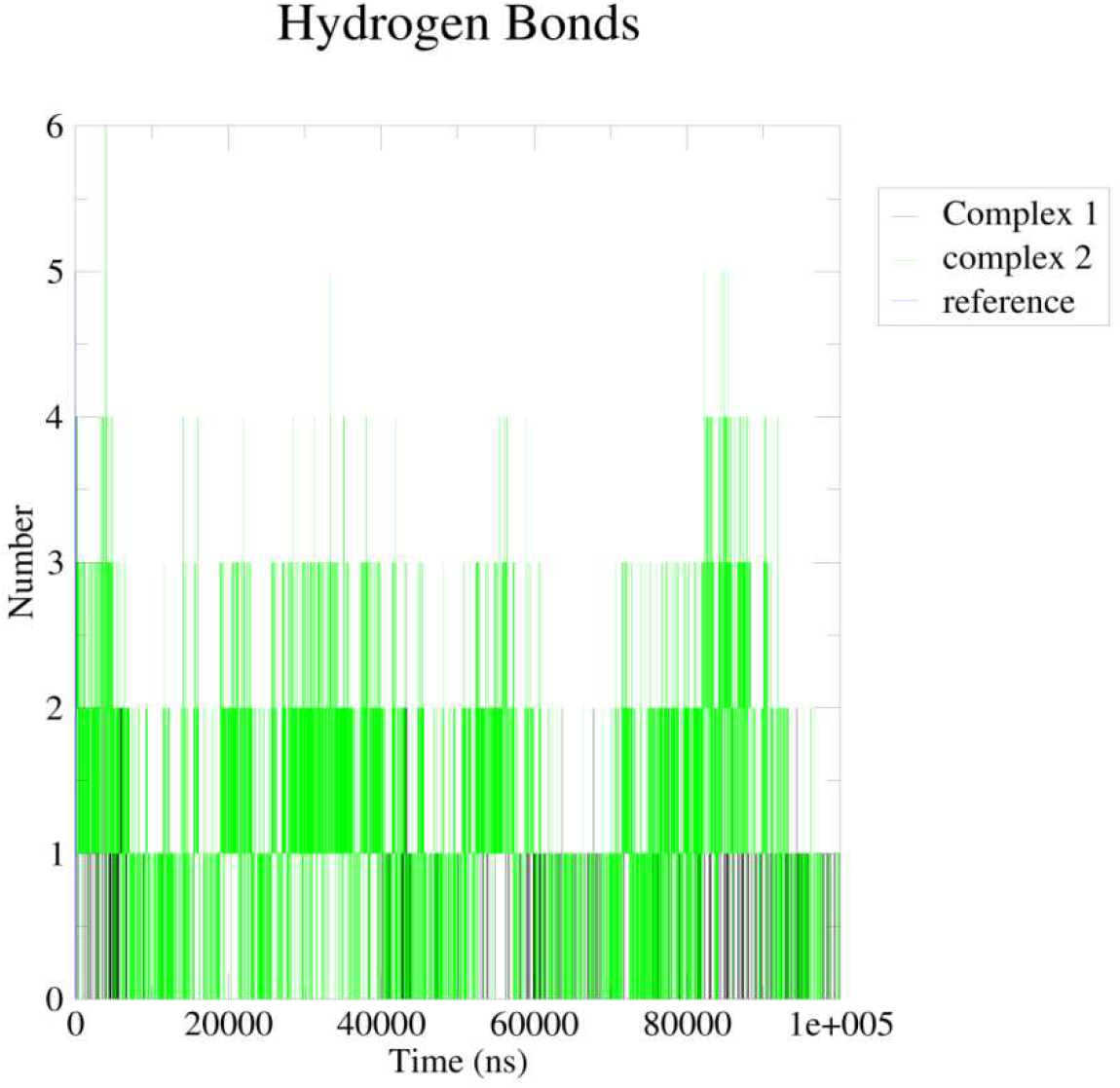
Hydrogen bond analysis for Reference compound, Complex 1,Complex 2

### 3.9 Principal component analysis

The principal contribution of the system’s overall dynamics is determined using the principal component analysis (PCA) technique.

Complex 1 has a cluster in the figure 1.9 which is black in colour and not has much deviation compared to complex 2.

**Fig 1.9:**
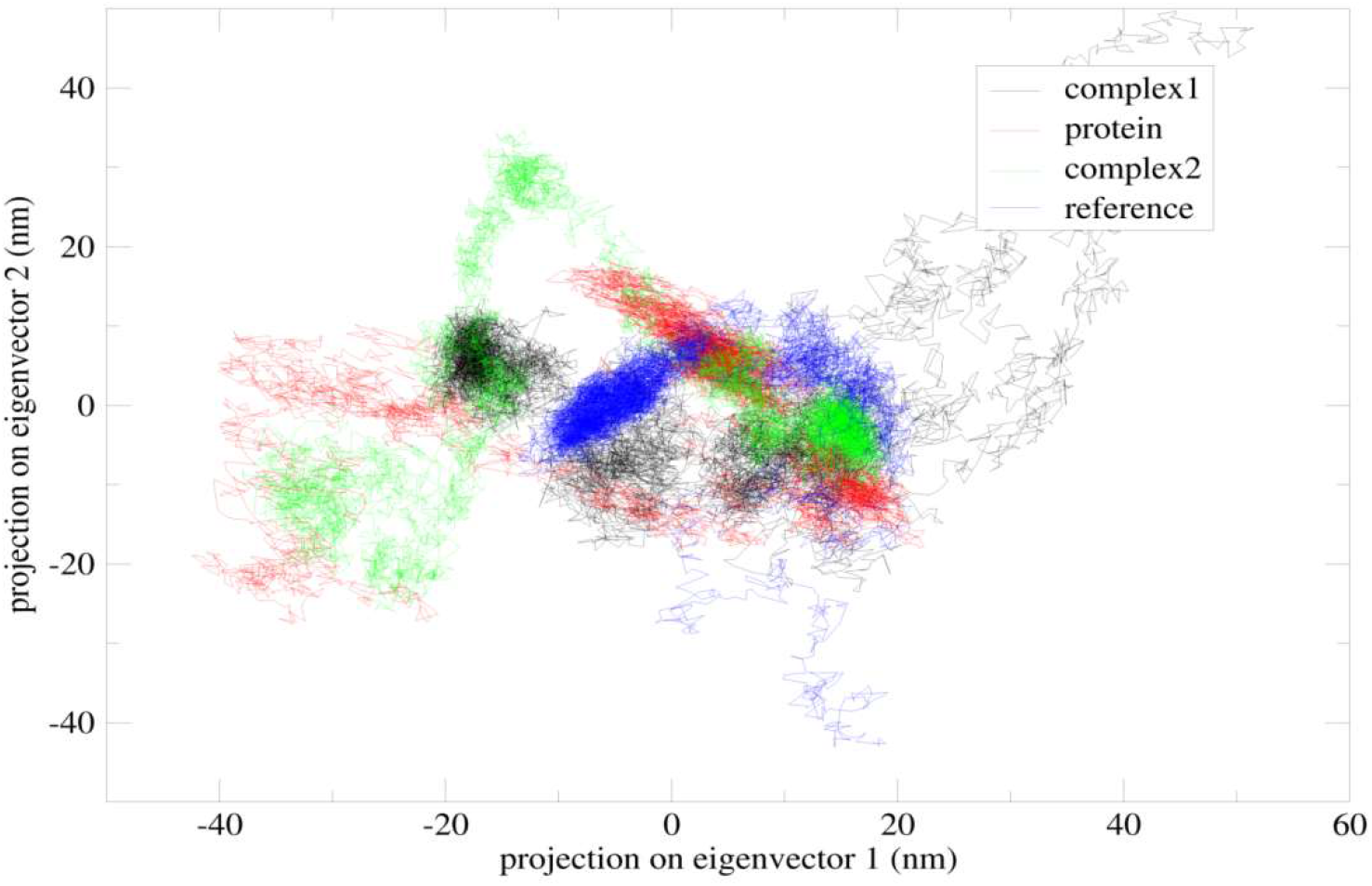
Principal component analysis for Reference compound, Complex 1,Complex 2.

### 3.10 Mmpbsa analysis

Chemical engineering Poisson-Boltzmann surface area (MM-PBSA), a method to determine the interaction free energy, has gained popularity in the study of biomolecular interactions.

## 4. Discussion

The ectodermal organs that make up teeth are hair, nails, and horn. Oral cancer is thought to be more common in India than anywhere else in the world, despite the fact that periodontal disease affects around 85% of the population and dental caries roughly 60% of Indians. An interplay between the epithelium and mesenchyme in a signal network controls tooth formation.

Supernumerary teeth, also referred to as teeth agenesis, are any teeth or orthodontic structures that are either more or fewer in number than the average person’s teeth. Supernumerary teeth arise as a result of USAG1 Protein dysfunction. Congenitally, between 0.1 percent and 3.6 percent of people have more or fewer teeth than they need. Dental impaction, delayed eruption, tooth misalignment, irregular tooth spacing, and follicular cyst development are among the complications connected to tooth agenesis.

USAG-1, a bifunctional protein mostly expressed in the kidneys, is known as the “uterine sensitization associated gene-1.” It regulates Wnt signaling and is a BMP (Bone Morphogenetic Protein) antagonist. Wnt signaling is crucial for the development of supernumerary teeth, whereas BMP signaling is crucial for the morphogenesis of additional teeth. Supernumerary teeth cannot form unless BMP signaling is inhibited. A C-terminal cysteine knot domain is present in the residues from 75 to 170 of the USAG1 protein (CTCK). As the protein’s preferred binding site, cysteine-104 in the USAG1 protein is primarily responsible for the protein’s dimerization. By inhibiting GSK-3, tideglusib boosts WNT signaling when applied to injured teeth. It leads to the repair of teeth.

USAG-1 was suggested as a potential therapeutic target for tooth regeneration by Suginami et al. They suggested using an anti-USAG-1 strategy to stimulate tooth regeneration, however it has been linked to a number of negative effects. A natural lead compound that can be used as a medicinal medication for tooth regeneration is the study’s major goal.

## 5. CONCLUSION

By modeling the protein, validating the modeled protein, screening of the natural molecules, ADMET prediction and the molecular dynamic simulation of the top two binders (3411,6473) and by the molecular dynamic simulation analysis, we can predict that 6473 is good non-toxic compound with greater stability and a good compactness compared to the other one can be considered for clinical trials and a promising drug target for teeth regeneration.

## Acknowledgement

We sincerely acknowledge the computation facility provided by TIFAC, SASTRA University

## Author’s declaration

KKG have designed the methodology. PR and EE have carried out the work.

